# Drug Repurposing through Meta-path Reconstruction using Relationship Embedding

**DOI:** 10.1101/2025.03.15.641108

**Authors:** Jiseong Park

## Abstract

De novo drug development are often costly, time-consuming and risky. Drug repurposing/repositioning, increasing usability of approved drugs, offers relatively high chance of success and efficiency based on the verified safety. Applying heterogeneous biological knowledge graph enabled compelling in silico drug repurposing, yet challenging to extract interaction between heterogeneous characteristics and utilizing multi-hop interactions. In this paper, I propose a PREDR model that predicts the drug-disease relationship by defining a drug-gene-disease meta-path from heterogeneous knowledge graphs and combining each heterogeneous relationship. The PREDR model reconstructs a meta-pathway based on relational information extracted by embedding each biological feature. The PREDR model outperformed compared to existing drug repurposing models in predicting drug-disease interaction, and demonstrated the effectiveness of meta-path reconstruction by showing higher performance than the result of learning and combining each heterogeneous relationships separately. The PREDR model can also explain the reaction mechanism of suggested drugs on the defined meta-pathway using heterogeneous interactions driven from reconstruction process. In predicting drug candidates for lymphoblastic leukemia disease conducted as a case study, the highest scored candidate is confirmed effectiveness of the disease in the literature, and predicted genes are verified to be targeted by both candidate and disease by various academic sources.

## 1 Introduction

### 1.1 Drug repurposing

De novo drug discovery and development takes an average of 13 years and 2-3 billion dollars that is too long period and expensive to practically available. The reasons of the time and cost expensive issue include the process comprising examination of efficacy, toxicity and pharmacokinetic and pharmacodynamic profiles for the target drug into cell- and animal-based studies and even to safety and efficacy in human subjects in clinical trials[1, 7, 8].

#### Advantages of drug repurposing

Drug repurposing (also called as drug repositioning) is a strategy for identifying new use for approved drugs which are already in use of their original indications[3]. Representative examples of drug repurposing include Viagra, which was recognized as an erectile dysfunction treatment in the existing angina pectoris drug[12], and Thalidomide, which was recognized as a multiple myeloma treatment in the morning sickness prevention drug[13]. Increasingly, researchers and clinicians are considering this strategy because of its strong positive effects. The advantages of this approach include the relatively short time from being approved and cost-efficient.

Moreover, the strategy studies drugs that are already known safety profiles that holds the potential to result in a less risky business.

### 1.2 Computational approaches for drug repurposing

#### 1.2.1 Diverse approaches for drug repurposing

The methodologies in the field of drug repurposing can be largely classified into in vivo, in vitro and in silico methods. In these methodologies, in vivo and in vitro, which exploit real biological phenomena, can be used for verifying real world outputs by utilizing real cell or tissue-based wet lab experiment or even with animal testing. In vivo approach is a study which the effects of various biological entities are tested on whole, living organisms such as animals, including human, and plants[15]. In vitro approach is a study which the biological tests are done in a laboratory environment using test tubes, petri dishes and so on with limited biological entities such as cells or tissues. Both methodologies have the advantage of being able to produce accurate results because they are tested using real biological objects[16]. However, both methods have critical disadvantages that the laboratory environment greatly affects the prediction accuracy, the process of confirming the interaction with tissues and organs is limited, and it is difficult to consider biological diversity and complexity. In silico approaches (also known as computational approaches) has various advantages, such as being able to simulate interactions between drugs and targets based on various data, requiring relatively little time and cost, and reducing the amount of animal testing[4]. In this sense, drug repositioning usually tends to proceed secondarily in vitro for drugs identified in silico.

#### 1.2.2 Diverse approaches for computational drug repurposing

There are three major methods of deug repurposing in an in silico, or computational manner, which are structure-based, ligand-based, and network-based. These three methodologies have their own advantages and drawbacks for some senses. The followings are the brief explanation for these methodologies.

##### Structure-based approach

The structure-based method simulates the three-dimensional information of the compound to simulate the reactivity. They use text or image based format for representing physical and chemical structure of entity. For example, FASTA[9] or SMILES[10] format data are key examples of structural data to be used for the methodology. However, it has the disadvantage that it is very difficult to apply in the case of compounds whose three-dimensional structure is not known or whose structure is difficult to understand[5].

##### Ligand-based approach

The ligand-based methodology is a simulation method based on the characteristics and interaction relationship between ligand and receptor, and has the advantage of being able to derive accurate results based on various physicochemical information. For example, QSAR[11] model can be characterized by a collection of well defined for exploring and exploiting ever growing collections of biologically active chemical compounds. However, it is difficult to collect data on ligand and target information and based on insufficient information can result inaccurate performance which is unreliable and that can be obsolete[5].

##### Network-based approach

Network-based methodology is a methodology widely used in the field of drug repurposing. This method is based on data in the form of a graph composed of nodes and edges. Network data is characterized by being able to cover not only the nature of each factor but also the relationship between factors. This has a great advantage as data for understanding biological characteristics made up of complex interactions of various factors[5, 6]. There are two types of networks; homogeneous and heterogeneous. Homogeneous network is a graphical representation which all types of nodes are the same. Heterogeneous networks, in contrast, includes nodes of different types are mixed. Here, the homogeneous network has the disadvantage that it is difficult to consider all biological entities. For this reason, heterogeneous network data that can consider various entities and their interactions is being used a lot. A typical example of a heterogeneous biological network is a knowledge graph, which was used to conduct this research.

### 1.3 Knowlegde graph based drug repurposing

#### 1.3.1 Biological knowledge graph

A biological knowledge graph is graph-type data that contains biological entities such as genes, proteins, diseases, and drugs and their relationship information[19]. For example, if you look at some connectivity of a specific knowledge graph, Wafarin, VKORC1 gene, and blood clot may be connected. This is because wafarin is an anticoagulant that inhibits the expression of the vitamin k epoxide reductase complex subunit 1, namely VKORC1 gene, and affects the activity of coagulation factors that require vitamin k, so it can be linked as an interacting factor on the knowledge graph[17, 18]. As such, the knowledge graph has the advantage of being very effective as data considering various biological factors and their interactions[20].

#### 1.3.2 Meta-path defined by knowledge graph

Another advantage of the knowledge graph is that it is easy to derive a meta-path. A meta-path is an ordered connection sequence of nodes and edges defined on a network that describes complex relationships between node types included in a graph[21]. For example, the following drug-gene-disease sequence can be defined as one meta-path. Reconstructing a graph based on a meta-path has the advantage of being easy to explain the mechanism of action between elements. In this study, I conducted a study using the drug-gene-disease meta-path.

#### 1.3.3 Multi-hop interaction calculation

The method of searching source-destination relation of multi-hop relation is quite straight forward. Given the interaction matrix for each hop-related interaction, we can easily obtain the multi-hop interaction counts by simply multiplying them[23]. Let 𝒯^*v*^ and 𝒯^*e*^ be a set of node types and edge types respectively. For a given heterogeneous graph *G* = (*V, E*), *f*_*v*_ : *V* →𝒯^*v*^ and *f*_*e*_ : *E* →*𝒯*^*e*^. In this graph, we can define meta-path *P* as follows.

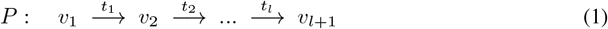

In the equation, *t*_*l*_ 𝒯∈^*e*^ and composite relation of the meta-path can be defined as *R* = *t*_1_ ○*t*_2_ ○… ○*t*_*l*_. Let *A*_*l*_ be an adjacency matrix for the edge type of *t*_*l*_, the adjacency matrix for the meta-path can now be calculated by multiplying those adjacency matrix like below.

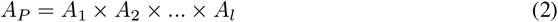

For example, looking at the graph in Figure 4, the graph consists of a combination of bipartite relations of blue, green, and orange node types. Here, when trying to understand the interaction from blue to orange, two intermediate bipartite relations, blue-green interaction and green-orange interaction, can be used. As a result of matrix multiplication of each binary interaction matrix, the blue-orange interaction matrix can be calculated. The elements of the matrix represent the path from blue to orange and the number of branches of the path. Therefore, in this way, multi-hop interactions can be calculated as a combination of bipartite relations or homogeneous relations of heterogeneous networks.

**Figure 1:**
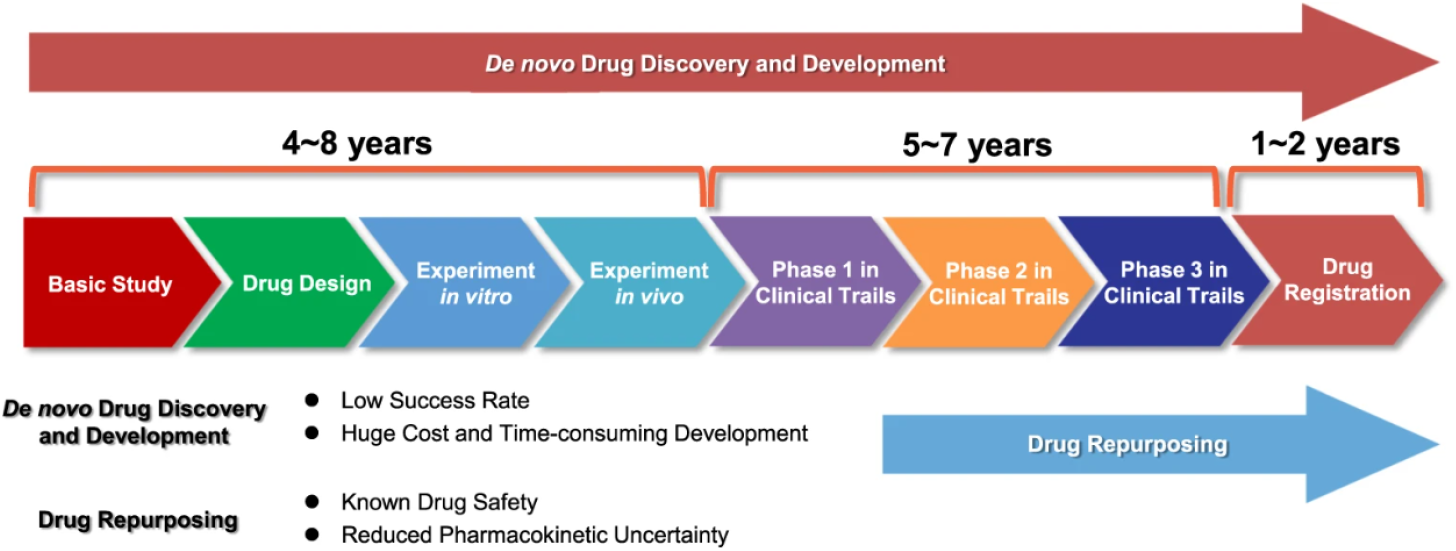
Drug repurposing takes relatively lower time and cost-efficient compared to de novo drug discovery and development.[2]

**Figure 2:**
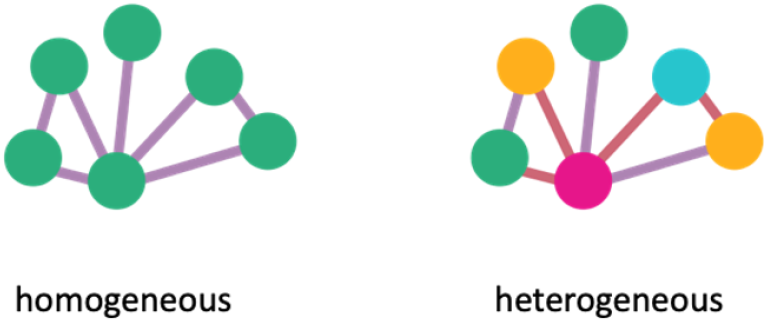
Example of homogeneous and heterogeneous graph.

**Figure 3:**
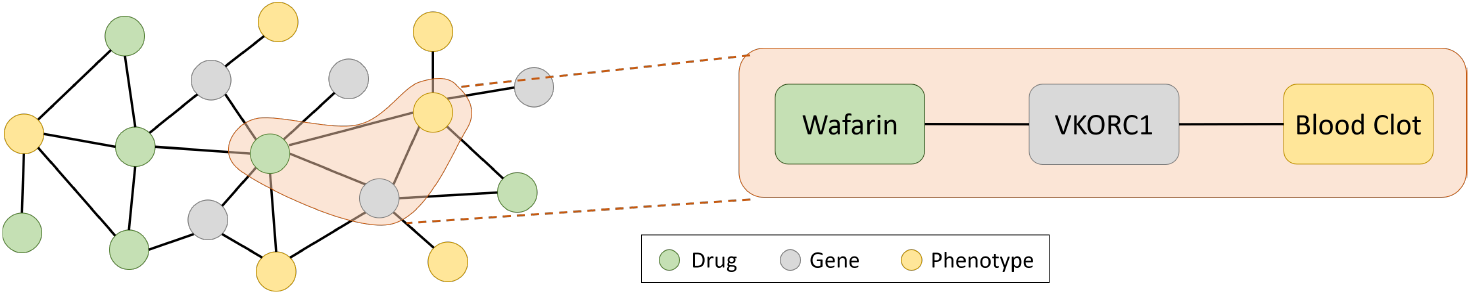
One example of biological knowledge graph and an example of its connectivity defined as drug-gene-phenotype.

**Figure 4:**
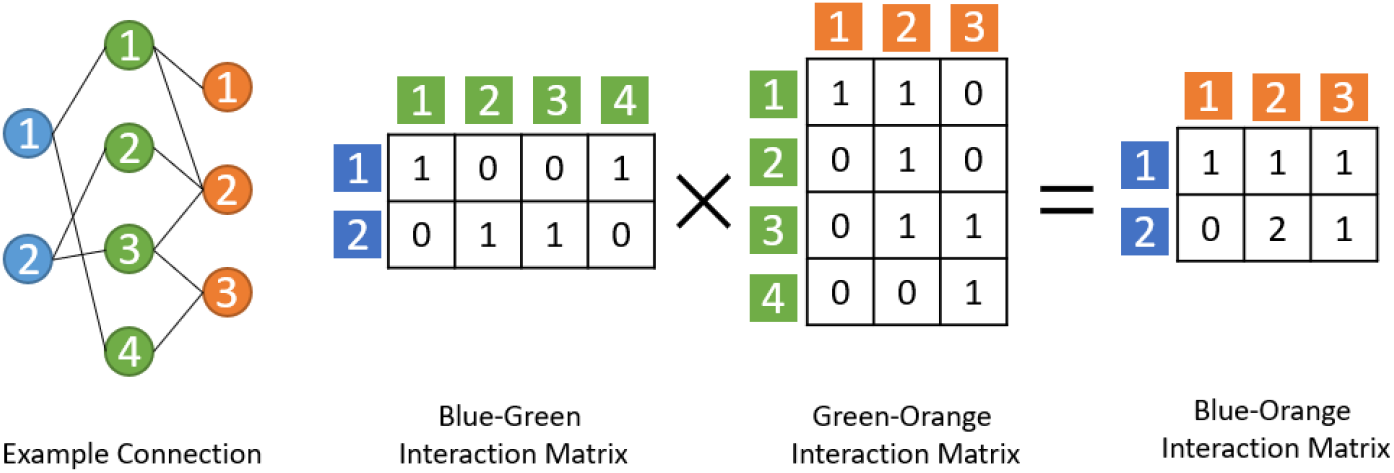
Brief illustration of how the multi-hop interaction can be considered using each intermediate relation.

#### 1.3.4 Meta-path based drug-target interaction prediction

The brief framework of utilizing meta-path to predict drug-target interaction is like Figure 5. As a basic structure, first collect various types of networks, build a knowledge graph based on them, define a meta-path, and extract various subgraphs based on the defined meta-path. Then, the model is used to predict the drug-target interaction by reconstructing the structure of the extracted graph[22].

**Figure 5:**
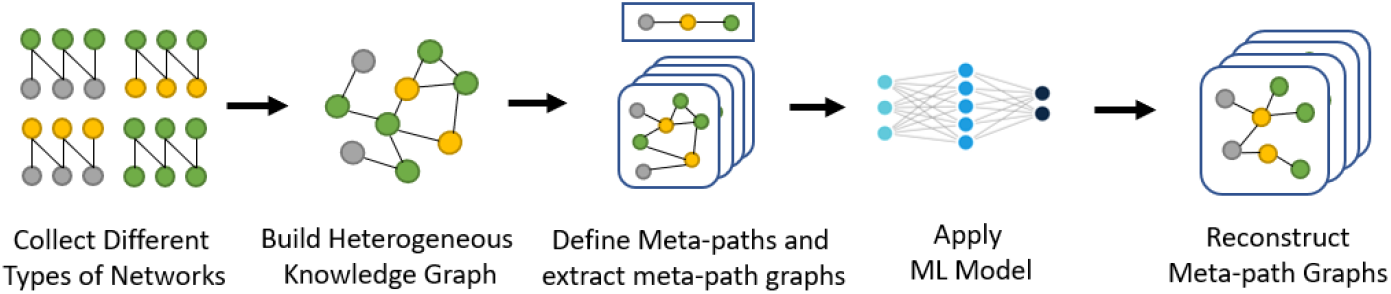
Conventional framework of knowledge graph based meta-path prediction.

**Figure 6:**
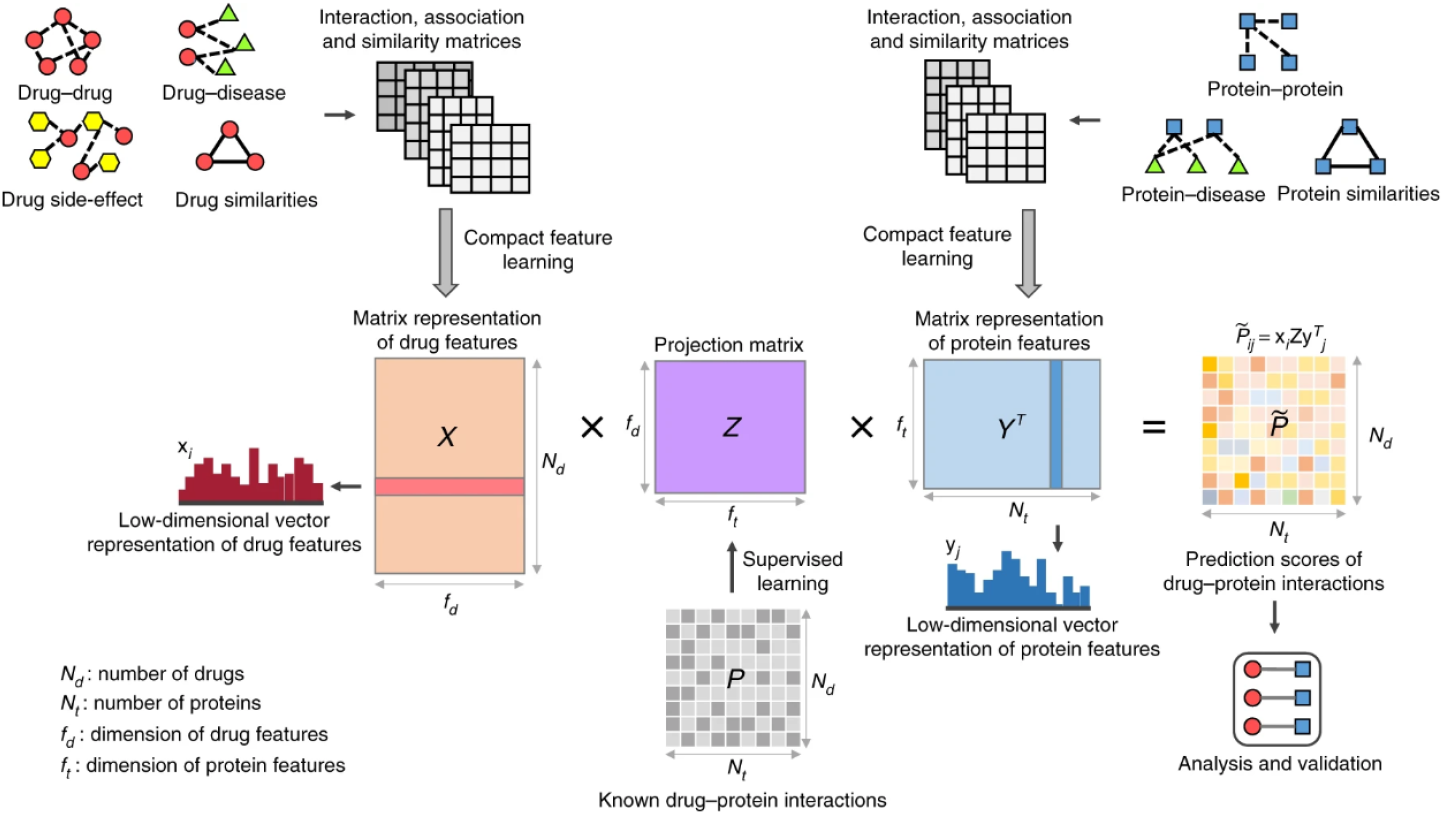
Method overview and framework for DTInet model.

**Figure 7:**
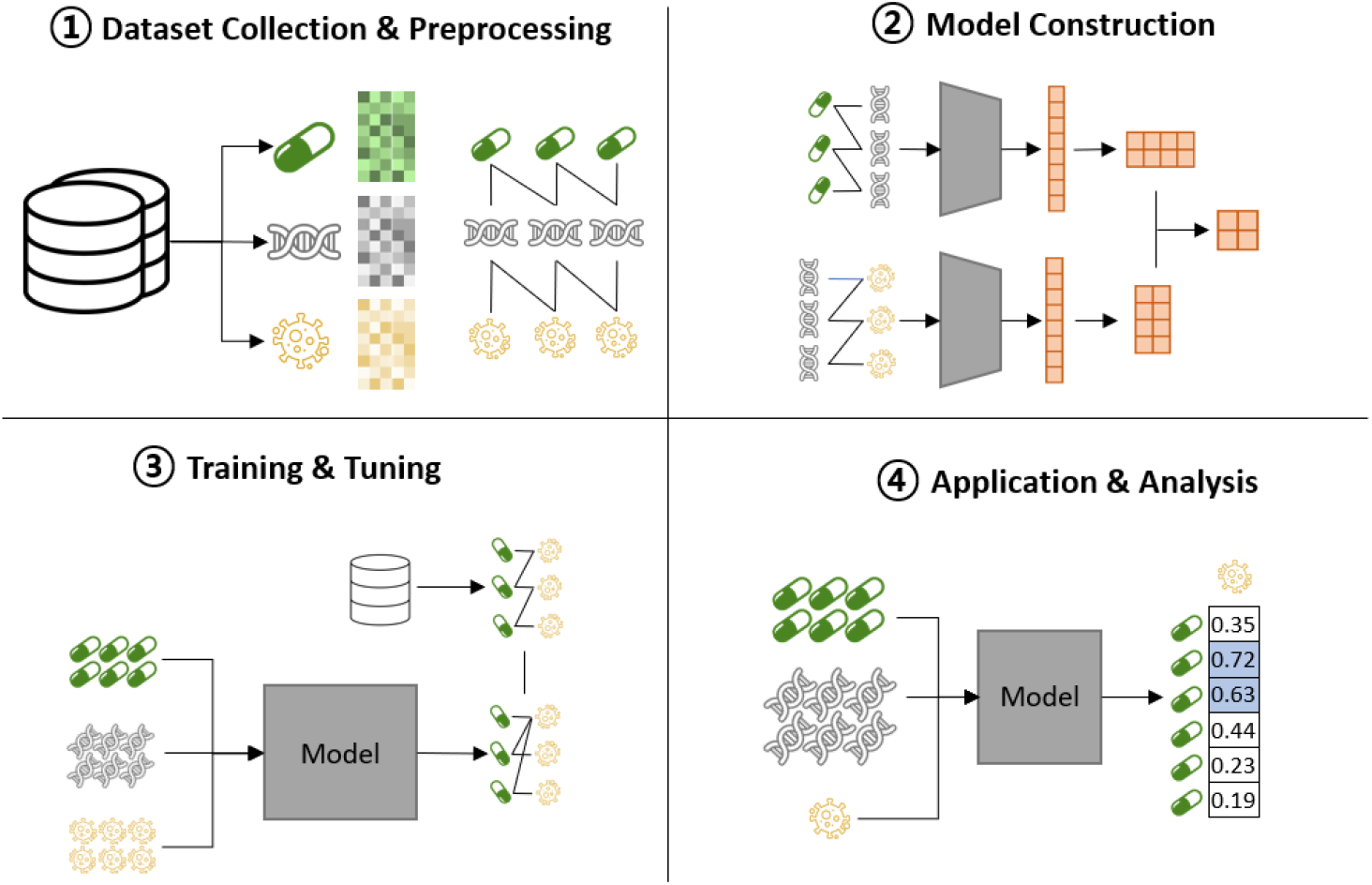
Method overview.

**Figure 8:**
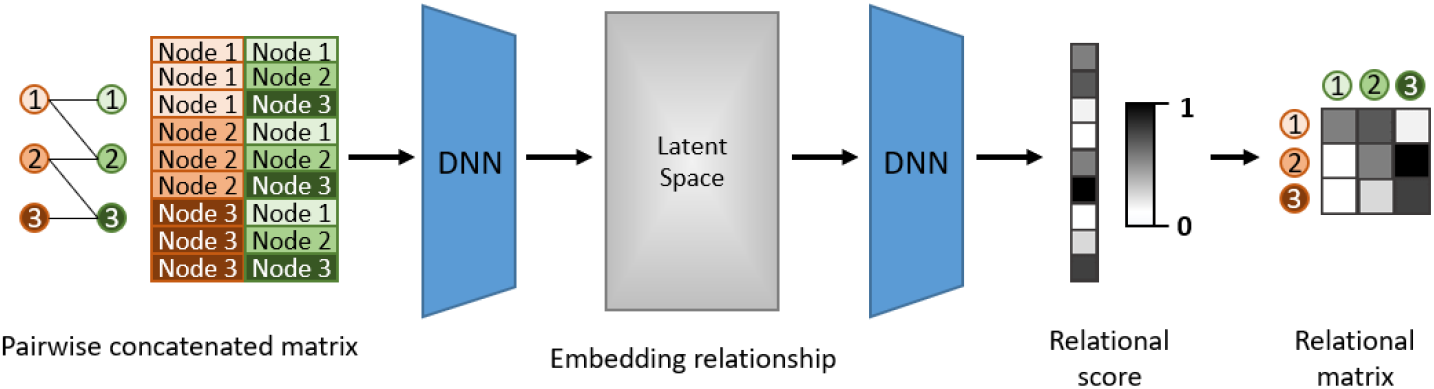
Bipartite relation embedding by pairwise concatenating each bipartite node feature vector.

### 1.4 Previous work

Researches on predicting drug-target interactions using heterogeneous knowledge graphs have been conducted in the past. Representatively, there is the DTINet[14] model introduced in the journal nature communications in 2017. The DTINet model was implemented to predict drug-target interactions by reconstructing the similarity between the two entities by introducing a projection matrix that learns the connection information to the cosine similarity of the feature vector of each biological entity. DTINet not only integrates diverse information from heterogeneous data sources (e.g., proteins, drugs, diseases and side-effects) but also copes with the noisy, incomplete and high-dimensional nature of large-scale biological data by learning low-dimensional but informative vector representations of features for both drugs and proteins. The key concept is similar from concept of other low-dimensional representation methods; using the low-dimensional feature vectors learned by DTINet capture the context information of indicidual networks, as well as the topoligical properties of nodes (e.g. drugs or proteins) across multiple networks. Based on these vectors, DTINet then finds an optimal projection from drug space onto target space, which enables the prediction of new DTIs according to the geometric proximity of the mapped vectors in a unified space.

However, these past methodologies have critical limitations that are highly rely on prior knowledge while extracting relationships. They also cannot explain the molecular mechanism of actions for newly discovered drugs.

### 1.5 Research objective

In this research, I will predict bipartite relation based on embedding pairwise concatenated feature vectors. Also, I will conduct the drug repurposing tasks to find some promising candidate for given specific disease and give insights of their mechanism by reconstructing meta-path for drug-disease relation through embedding bipartite relation.

## 2 Method

### 2.1 Method overview

In order to conduct the research, three biological entities, which are drug, gene and disease, and their heterogeneous relations are collected from various open accessible databases. These collected raw data are then preprocessed that can be fit to the suggested model and other baseline models. Next, defined and construct the model which can reconstruct the predefined meta-path, and then be trained and tuned using the preprocessed data. Afterward, based on the completed model, drug-disease interaction prediction tasks are conducted. We also run a practical drug repurposing exercise including extracting the drug candidates for the specific disease and analyzing the molecular mechanisms based on the designated meta-path.

### 2.2 Dataset statistics

In this research, three biological entities are collected for the dataset: drug, gene and disease. The drug was used by reducing the length of 1024 ECFP vector to 5 dimensions in the DrugBank[24] database. Genes were expressed in a 5-dimensional vector by integrating how target-specific the gene was in the DisGeNet[25] database, how extensive the mutation was, how resistant it was to mutation, and what kind of protein the gene was expressed in. For the disease, a 5-dimensional vector was constructed based on the information that could be extracted from the BioSNAP[26] database on the disease-disease homogeneous network.

For each relationship information, drug-gene relationship information and gene-disease relationship information to be used as heterogeneous relationship inputs were brought from the BioSNAP[26] database, and information on actual drug use to be used as a label was brought from the DrugBank[24] database.

Based on the collected data, I constructed subgraphs with various combinations of drugs, genes and diseases introduced at Table 1. Each subgraph *G* consists of 3 feature matrices; drug, gene and disease and label matrix, that is *G* = (*X*_*Dr*_, *X*_*G*_, *X*_*Dis*_, *A*_*label*_) where 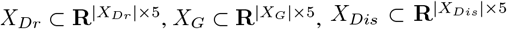 and 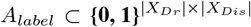 where |*X*_*Dr*_|, |*X*_*G*_|, |*X*_*Dis*_| are the number of rows for *X*_*Dr*_, *X*_*G*_, *X*_*Dis*_ respectively. Based on the subgraphs, I constructed a total of 4045 subgraphs to form a dataset for the model, that is, *Dataset* = {*G*_1_, *G*_2_, …, *G*_4045_}.

**Table 1:** Statistics of biological entities.

| Entity | Feature (n=5) | Volume | Database |
| --- | --- | --- | --- |
| Drug | ECFP (n=1024) | 4,045 | DrugBank |
| Gene | DSI, DPI, PLI, Protein Categories | 7,964 | DisGeNet |
| Disease | Disease-disease network based components | 519 | BioSNAP |

**Table 2:** Statistics of biological relations.

| Relationship | Purpose | Volume | Database |
| --- | --- | --- | --- |
| Drug-Gene | Intermediate relation | 15,307 | BioSNAP |
| Gene-Disease | Intermediate relation | 21,357 | BioSNAP |
| Drug-Disease | Approved indication | 95,621 | DrugBank |

### 2.3 Bipartite relationship reconstruction module

Bipartite relation embedding is a key concept to extract relation information inference strategy used in this paper. Based on the nodes of the provided graph, a pairwise concatenated matrix was created by representing all combinations of each node as a combination of feature vectors. Pairwise concatenated matrices are represented by all combinatorial pairs of nodes, so the number of rows in the matrix will be the multiplication of total number of nodes of each type. The pairwise concatenated matrix formed in this way was embedded using a multi-layer neural network(MLP), and relational information was reconstructed based on the embedded space.

The detailed method of constructing pairwise concatenated matrix is like follow. Let the bipartite graph *G* = (*N*_1_, *N*_2_, *V*) where *N*_1_ and *N*_2_ are each type of node set for the bipartite graph, that is, *N*_1_ = {*n*_11_, *n*_12_, …, *n*_1*k*_} and *N*_2_ = {*n*_21_, *n*_22_, …, *n*_2*l*_} which size of *k* and *l* respectively and *V* is a set of relation between *N*_1_ and *N*_2_. Also, the embedding dimension for each set of node, *N*_1_ ⊂ **R**^*m*^ and *N*_1_ ⊂ **R**^*n*^ that is, each node for each type of node has its embedding dimension of *m* and *n* respectively. In this sense, the pairwise concatenation matrix(PCW) can be constructed as shown in Equation 3.

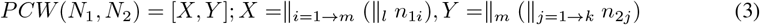

In the Equation 3, *X*⊂ **R**^(*k*×*l*)×(*m*+*n*)^ be a rowwise concatenated matrix for each node in *N*_1_ tiled for |*N*_2_| times where |*N*_2_| indicates the number of elements in *N*_2_ and *Y*⊂ **R**^(*k*×*l*)×(*m*+*n*)^ be a rowwise concatenated matrix for the rowwise concatenated matrix of *N*_2_ repeated for |*N*_1_| times where |*N*_1_| indicates the number of elements in *N*_1_, and∥ · indicates rowwise concatenation operation.

The constructed pairwise concatenated matrix then be passed through simple MLP models that can produce final relational vector **r** ∈ **R**^(*k*×*l*)^. This vector represents the probability for each bipartite node relation. The relation vector **r** can be reshaped into (*k* × *l*) form which can be interpreted as the bipartite adjacency matrix format *A*^′^ ⊂ **R**^*k*×*l*^. The bipartite adjacency matrix also can be used as an intermediate interaction matrix to obtain final meta-path relation by multiplying diverse intermediate interaction matrices introduced in Section 1.3.

### 2.4 Meta-path reconstruction model

In this paper, I suggest the model named PREDR(Pairwise Relation Embedding for Drug Repurposing) which aims to reconstruct meta-path by embedding each heterogeneous bipartite relation using pairwise relation embedding for drug repurposing. The Figure 9 shows the diagram of PREDR model architecture. Since I use the meta-path of drug-gene-disease, I created a reconstruction module that embeds drug-gene relationship information and gene-disease relationship information, respectively, and proceeded with end-to-end learning.

**Figure 9:**
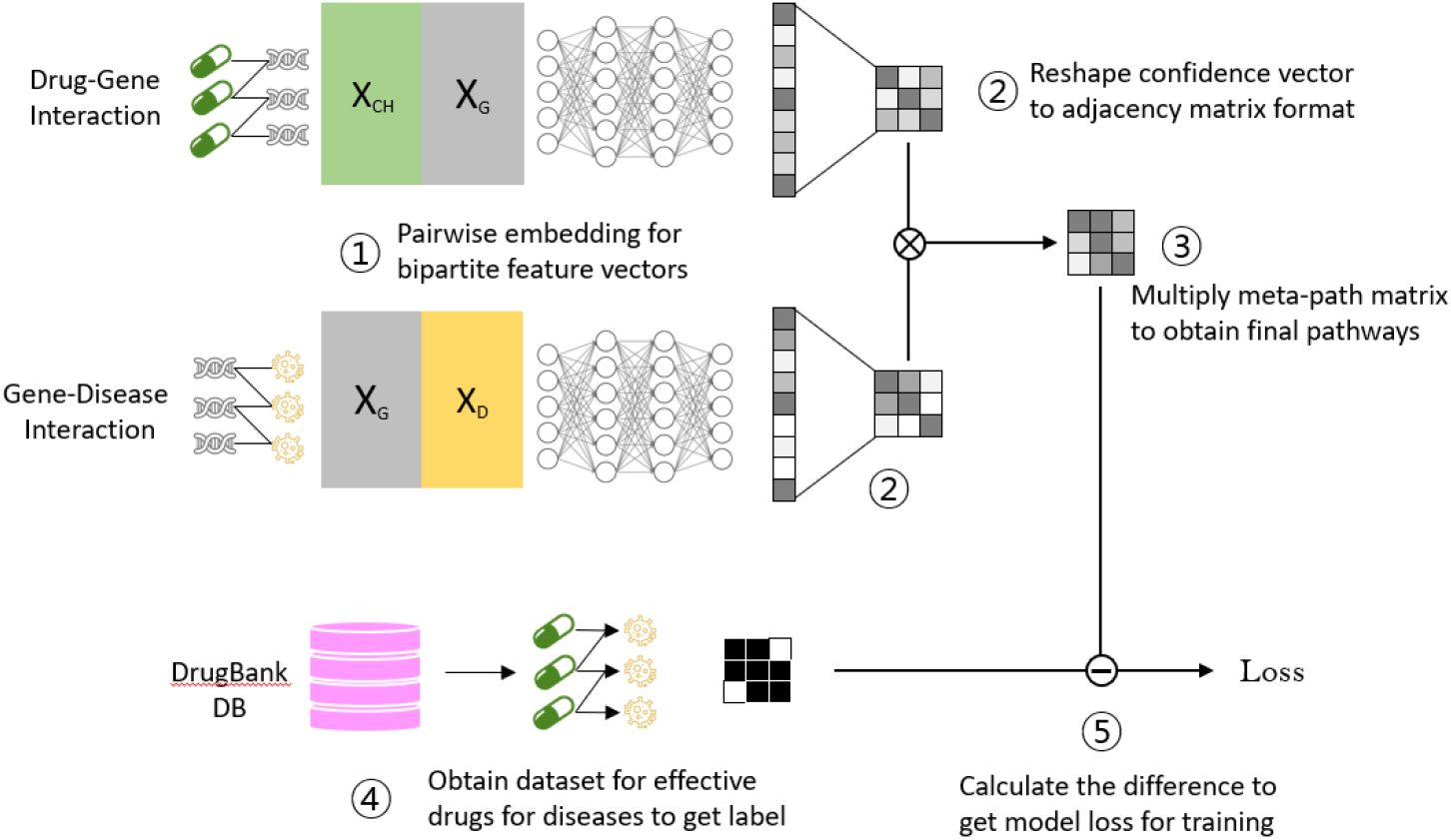
PREDR model architecture.

**Figure 10:**
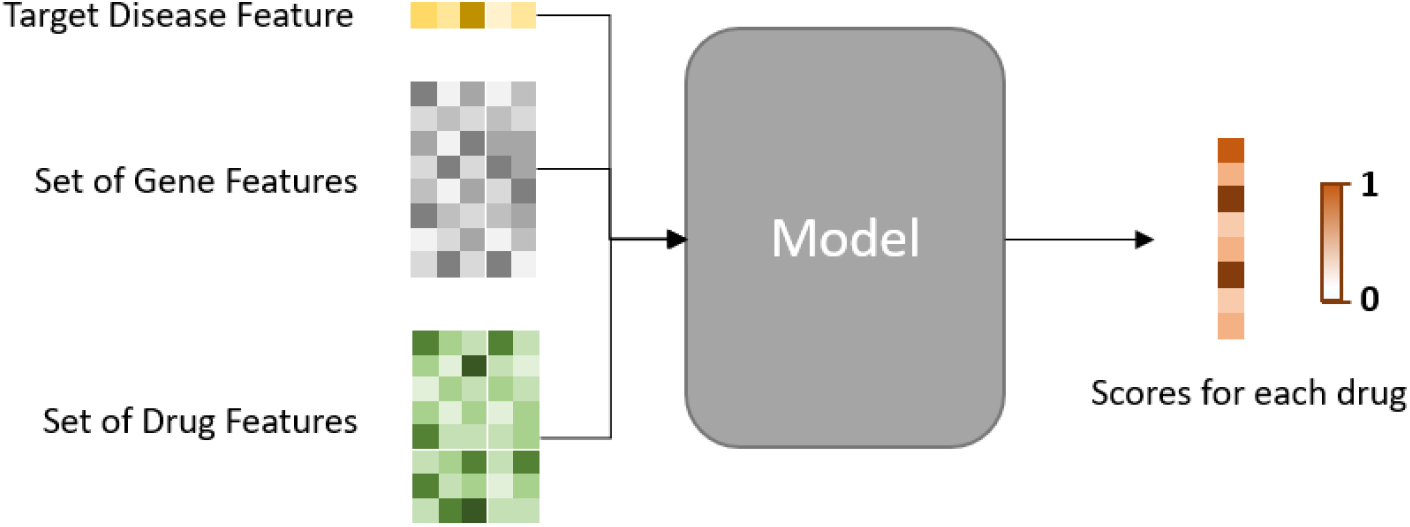
PREDR model utilization to obtain drug candidates for given disease.

**Figure 11:**
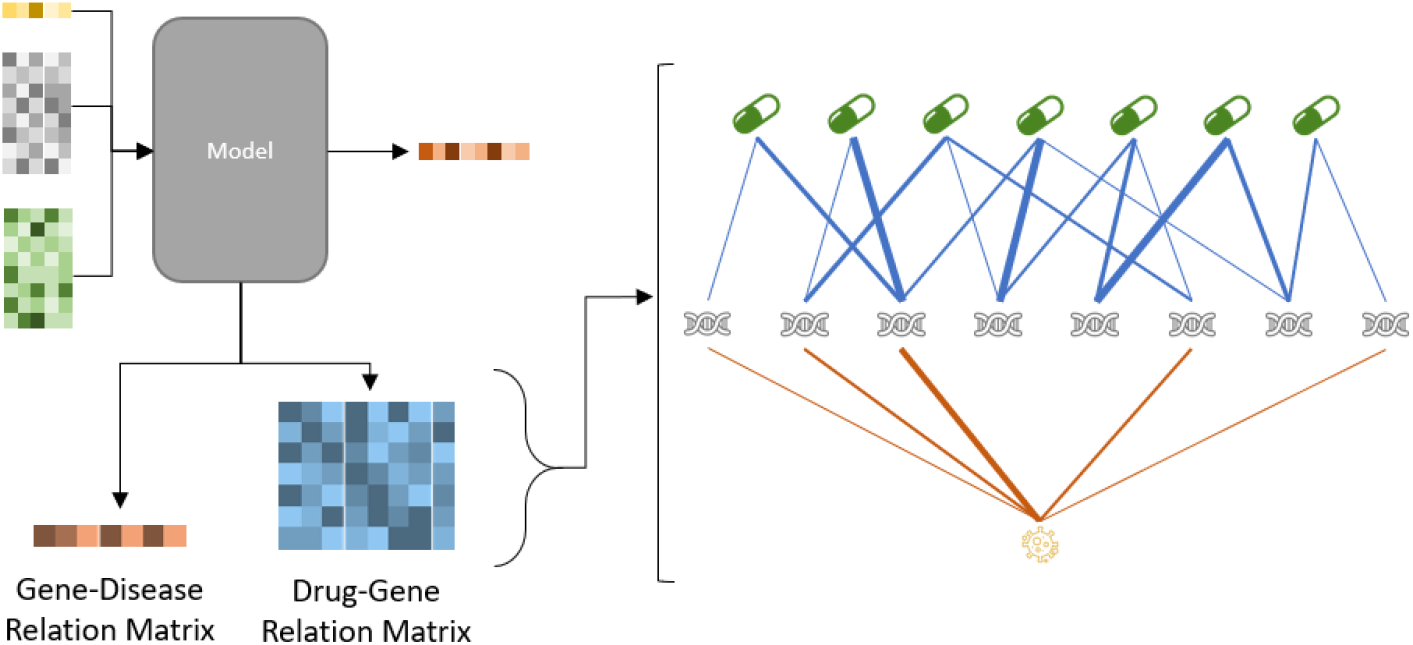
By the drug-gene matrix and gene-disease matrix inside, the model can be used to interpret the mechanism of action for suggested drug-disease relations.

First, I provided the subgraphs consists of drug, gene and disease feature matrices as inputs for the model and label matrix as a label for the input. In a subgraph, each feature matrix *X*_*Dr*_, *X*_*G*_, *X*_*Dis*_ for drug, gene and disease respectively, have their own variable number of rows. The feature matrices then be passed through 2 different bipartite relationship reconstruction modules; drug-gene and gene-disease. First, the drug feature matrix *X*_*Dr*_ and gene feature matrix *X*_*G*_ go into the drug-gene bipartite relation reconstruction module. The inputs then be constructed as a pairwise concatenated matrix *H*_*Dr*− *G*_(4), and embedded through message passing embedding network(MP), and then the reconstructed relationship vector (**r**_*Dr*− *G*_) is converted into a matrix based on the number of drugs and genes provided to form a drug-gene relationship matrix 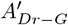(5 - 6).

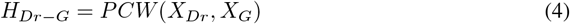

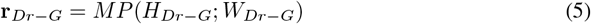

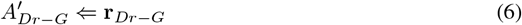

In the same way, the gene feature matrix *X*_*G*_ and the disease feature matrix *X*_*Dis*_ go into the gene-disease bipartite relation reconstruction module. The inputs then be constructed a pairwise concatenated matrix *H*_*G*−*Dis*_(7), embedded through message passing embedding network(MP), and converted to the reconstructed relationship information vector (**r**_*G*−*Dis*_) and also the matrix form 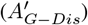 based on the number of genes and diseases provided(8 - 9).

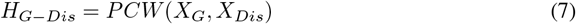

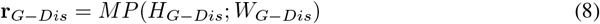

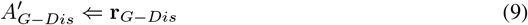

In these two module, each MP which embed the pairwise concatenated matrix for each bipartite relation is designed by using multi-layer perceptron(MLP) model. The framework of the module includes 3-layered neural network that consists of 2 layers for embedding input into smaller dimension and the other for shrinking the latent space into vector format. The structure of the embedding layer is like Equation 10.

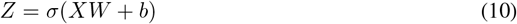

In this equation, *Z* represents a latent space for input matrix *X, W* and *b* represents learning parameters and bias. The activation function of the equation is designed to rectified linear unit(ReLU) which conserves the positive elements and cancels the negative elements to zero, that is,

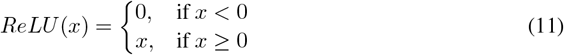

In this model, the original input matrix *X* will have its dimension of 10 since the matrix is formed by pairwise concatenation of two feature vectors of dimension 5 for each. By passing two embedding layers, the input matrix is mapped into lower dimensional space. As a result of hyperparameter tuning, the dimension of mapping space is designated to 8 and 4 respectively. Now, the final latent space then be shrank into vector shape to form a relationship vector. The structure of the relation reconstruction layer is like Equation 12.

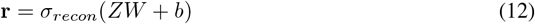

In this equation, *σ*_*recon*_ is exponential linear function(ELU) which shape is defined as Equation 13.

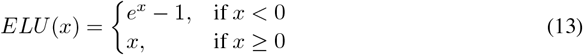

By multiplying the two relationship metrics 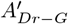 and 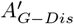 obtained by the two modules in this way, a drug-disease relationship matrix 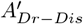 that can infer the possibility of a drug-gene-disease meta-path is finally calculated (14).

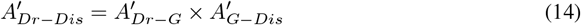

The loss of this model is calculated using the actual use of drug. The actual drug utilization was extracted from the DrugBank database. Each label was constructed as drug-disease interaction matrix format *A*_*label*_ corresponding to the input drugs and diseases. The matrix is symmetric and binary, that is, 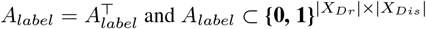 where |*X*_*Dr*_| and |*X*_*Dis*_| are the number of rows for *X*_*Dr*_ and *X*_*Dis*_ respectively which mean the number of input drugs and diseases. Now, the loss of the model is then be calculated by the binary crossentropy of each component of label matrix and model output as like below.

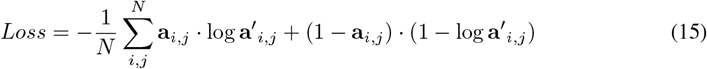

In the Equation 15, **a**_*i,j*_ and **a**^′^_*i,j*_ indicate element of *i*^*th*^ row and *j*^*th*^ column of *A*_*label*_ and 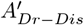 respectively and N denotes the total number of elements of *A*_*label*_. Using the loss calculated, the model learning was conducted by back-propagation with it.

### 2.5 Model application

#### 2.5.1 Application of the trained model for drug repurposing

When the disease target for repurpose and the characteristic information of various genes and drugs are provided as inputs to the trained model, a score vector for each drug’s association with the disease is calculated. Based on this vector, it is possible to identify which drugs are suitable for use in this disease, and based on this, candidates for drug repositioning can be created.

#### 2.5.2 Interpretation of mechanisms for the candidates

In the model structure described above, drug-gene relationship matrix and gene-disease relationship matrix can be provided as intermediate outputs. The matrix are matrix that contain drug-gene relationship or gene-disease relationship, respectively, and drug-gene-disease pathways can be analyzed based on the matrix. Based on these analyzed pathways, it is possible to gain insight into how the repurposed drug might work against the actual disease.

## 3 Result

### 3.1 Overall Performance of Drug-Disease Interaction

#### 3.1.1 Baseline models

We compare the performance of our model against various machine learning models that can be applied to drug repurposing task. Specifically, we compare our model’s performance against DTINet[14], and two basic machine learning classifiers which are multi-layer perceptron (MLP)[27] and support vector machine (SVM)[28].

**DTINet** Basically, the DTINet model is a model designed for drug target interactions. However, the study explains that the proposed model can contribute to drug repurposing by predicting drug disease interactions. Therefore, in the drug and target (protein) provided as the input of the model, the target data was replaced with the disease data. The provided input feature vector used the same feature vector as the dataset created in this study.

**MLP** The MLP model consisted of a total of three hidden layers, and the embedding dimensions were set to 16, 8, and 4, respectively. Activation used rectified linear unit (ReLU). The learning rate was set to 0.001, the momentum was set to 0.9, and the ADAM optimizer was used. A total of 200 epochs were trained without learning rate decay and early stopping.

**SVM** The SVM model used the linear support vector classifier model provided by the scikit-learn python package. The intercept scaling was set to 1, and a total of 1000 iterations were repeated and trained. In addition, the random state value was provided as the same value for fair performance comparison with the MLP model.

#### 3.1.2 Validation dataset preparation

From the features and relations retrieved from various databases mentioned at Section 2.2, we collected 4045 subgraphs considering drug-gene-disease meta-path. The obtained dataset than be separated into 3 groups; training set, validation set and test set. In this dataset, each data consist of 6 parts; feature matrix of drug set, feature matrix of gene set, feature matrix of disease set, drug-gene heterogeneous adjacency matrix, gene-disease heterogeneous adjacency matrix and drug-disease heterogeneous adjacency matrix.

#### 3.1.3 Validation Metrices

In this paper, I used three major metrices for assessing model performance; accuracy, auroc, and aupr. According to the confusion matrix, the accuracy can be defined as Equation 16. The AUROC also known as AUC - ROC curve is a performance measurement for the classification problems at various threshold settings. ROC is a probability curve and AUC represents the degree or measure of separability. It tells how much the model is capable of distinguishing between classes. Higher the AUC, the better the model is at predicting 0 classes as 0 and 1 classes as 1. The AUROC curve is plotted with TPR(eq. 18) against the FPR(eq. 19) where TPR is on the y-axis and FPR is on the x-axis. The area under the precision-recall curve (AUPRC) is a useful performance metric for imbalanced data in a problem setting where you care a lot about finding the positive examples. The AUPRC is calculated as the area under the PR curve. A PR curve shows the trade-off between precision(eq. 17) and recall(eq. 18) across different decision thresholds.

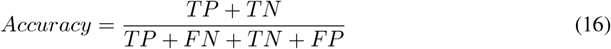

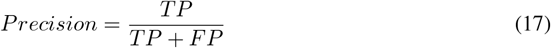

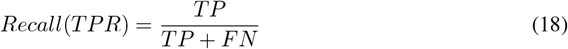

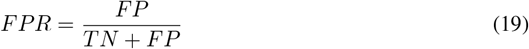

#### 3.1.4 Drug-disease interaction prediction performance

Learning was conducted through 5-fold cross validation[29], and the dataset provided for each training provided the ratio of positive and negative labels at 1:1. The Figure 12(a) is data showing the results of learning in terms of three evaluation indicators mentioned in advance. As a result of comparing the performance of the existing DTInet model, the simple MLP model, and the SVM model, it was confirmed that the performance of the PREDR model was comparable to other models. Since the ratio of positive label and negative label was set to 1:1, AUROC was determined to be the most effective evaluation index among the three evaluation metrices mentioned prior. As of this concern, an additional analysis of the AUROC score was conducted for the performance of each model. As a result shown in Figure 12(b), in the performance distribution derived as a result of 5-fold cross validation, PREDR proved to outperform the existing model, DTINet, by showing a performance distribution with a p-value of less than 0.05, and for baseline models such as SVM and MLP, the p-value of less than 0.01. Therefore, it showed a performance distribution differences were quite significant so that it proved that it worked well for all baseline models.

**Figure 12:**
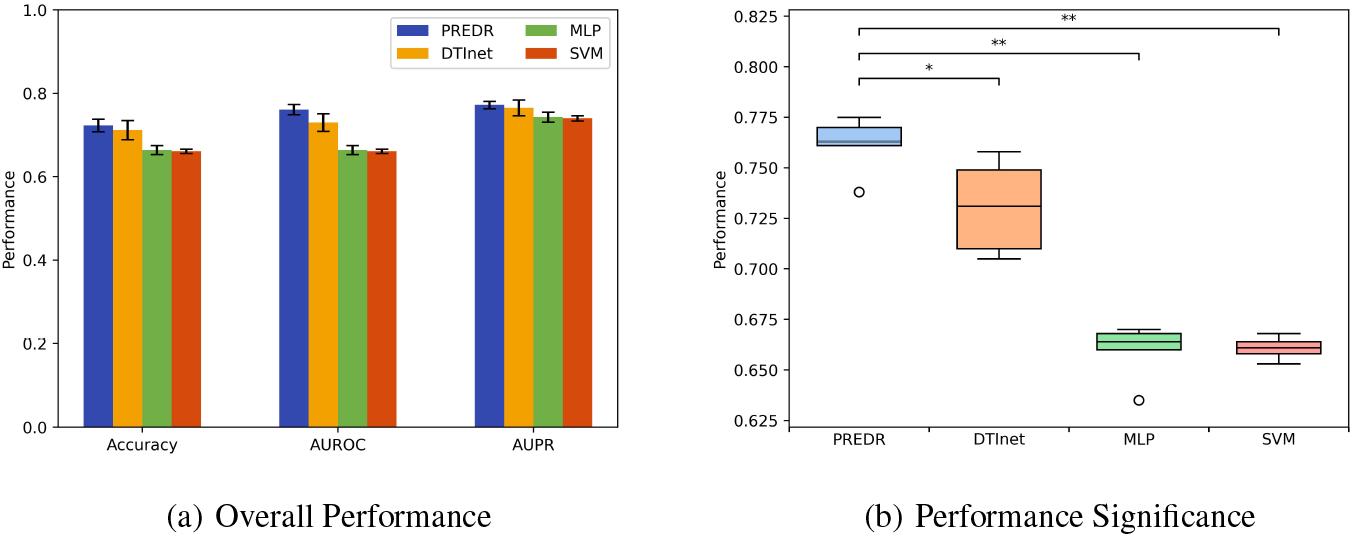
(a) Performance of PREDR model compared to baseline models. (b) AUROC performance of PREDR model compared to baseline models. According to the significance, PREDR outperforms to baseline models.

**Figure 13:**
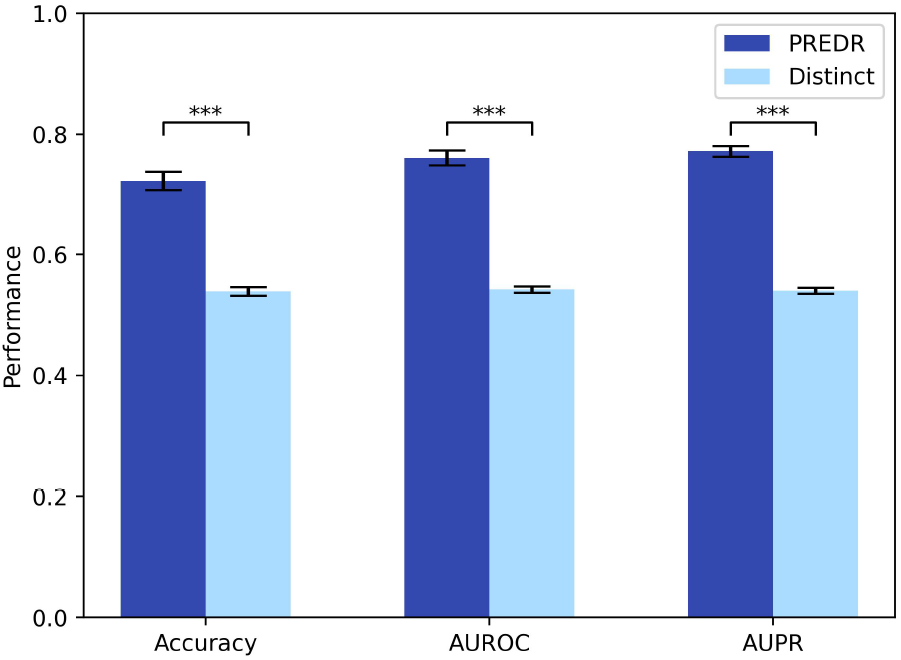
Performance of PREDR model compared to modified strategy conjugate separately trained modules.

### 3.2 Efficacy of Meta-path Reconstruction

This is a comparison of the performance of the PREDR model with that of reconstructing after training each module individually without connecting each module end-to-end. The chart compares the performance of the reconstructed meta-path by multiplying the results of each separately learned module and the reconstructed result through PREDR in three evaluation indices: accuracy, auroc, and aupr. It was confirmed that each module showed satisfactory performance in reconstructing heterogeneous relationships, but based on the results of learning separately, it was confirmed that the performance was much better when the model was constructed end-to-end than when the meta-path reconstruction was performed.

### 3.3 Case Study

#### 3.3.1 Lymphoblastic leukemia

Acute lymphoblastic leukemia and lymphoblastic lymphoma constitute a family of genetically heterogeneous lymphoid neoplasms derived from B- and T-lymphoid progenitors.[31] If you have leukaemia your body makes some abnormal blood cells[30]. These leukaemia cells behave differently from healthy blood cells. Acute means that it develops quickly and needs to be treated straightaway. Diagnosis is based on morphologic, immunophenotypic, and genetic features that allow differentiation from normal progenitors and other hematopoietic and nonhematopoietic neoplasms. ALL is rare. Around 800 people in the UK are diagnosed with ALL each year. It can develop at any age. However, most ALL cases occur in children, with an incidence of 3 to 4/100,000 in patients 0 to 14 years of age and >1/100,000 in patients older than 15 years, in the United States.[34] The prognosis of ALL has improved dramatically over the past several decades as a result of adapting therapy to the level of risk for relapse, improvements in supportive care, and optimization of the existing chemotherapy drugs.

#### 3.3.2 Recall for the original indications

In order to conduct a case study through the trained model, a feature vector for lymphoblastic leukemia was first prepared and provided as an input to the model. In addition, a set of drugs and a set of genes were prepared from the prepared dataset to form feature matrices, respectively, and provided as inputs to the model. A total of 29 drugs were used, and 15 of them were drugs whose usability had not been verified for lymphoblastic leukemia. Also, the number of genes used was 141. All of the drug, gene, and disease features used at this time were not used in the training stage. Using this, the model created a drug-gene interaction matrix of 29 × 141 and a gene-disease interaction matrix of 141× 1 as intermediate interaction matrices, and finally created score vector for 29 drugs for lymphoblastic leukemia.

The drugs listed in the Table 4 are drugs published in DrugBank database for which already have been approved for use in lymphoblastic leukemia. The PREDR model is a model for binary classification problems, and drugs with a final score of 0.5 or higher are classified as having utility. Note that, as described on the Figure 14 the output score for each drug has maximum of 0.590 and minimum of 0.395. That is, the classification problem for the lymphoblastic leukemia was quite challenging compared to the other diseases. However, PREDR determined that 11 of these 14 drugs were available, resulting in a recall of about 0.79 for the disease.

**Table 3:** Performance of each separately trained drug-gene and disease-gene relation reconstruction modules.

|  | Drug-Gene | Disease-Gene |
| --- | --- | --- |
| Accuracy | 0.68 | 0.75 |
| AUROC | 0.72 | 0.82 |
| AUPR | 0.71 | 0.81 |

**Table 4:** The 14 drugs are indicated to be used for treating lymphoblastic leukemia and 11 out of 14 drugs are verified as available by the PREDR.

| No. | Drug name | Score |
| --- | --- | --- |
| 1 | Nelarabine | 0.568 |
| 2 | Daunorubicin | 0.562 |
| 3 | Prednisone | 0.556 |
| 4 | Doxorubicin | 0.552 |
| 5 | Dasatinib | 0.550 |
| 6 | Vincristine | 0.539 |
| 7 | Imatinib | 0.533 |
| 8 | Dexamethasone | 0.531 |
| 9 | Cyclophosphamide | 0.514 |
| 10 | Ponatinib | 0.509 |
| 11 | Methotrexate | 0.501 |
| 12 | Clofarabine | 0.496 |
| 13 | Mercaptopurine | 0.490 |
| 14 | Cytarabin | 0.465 |

**Figure 14:**
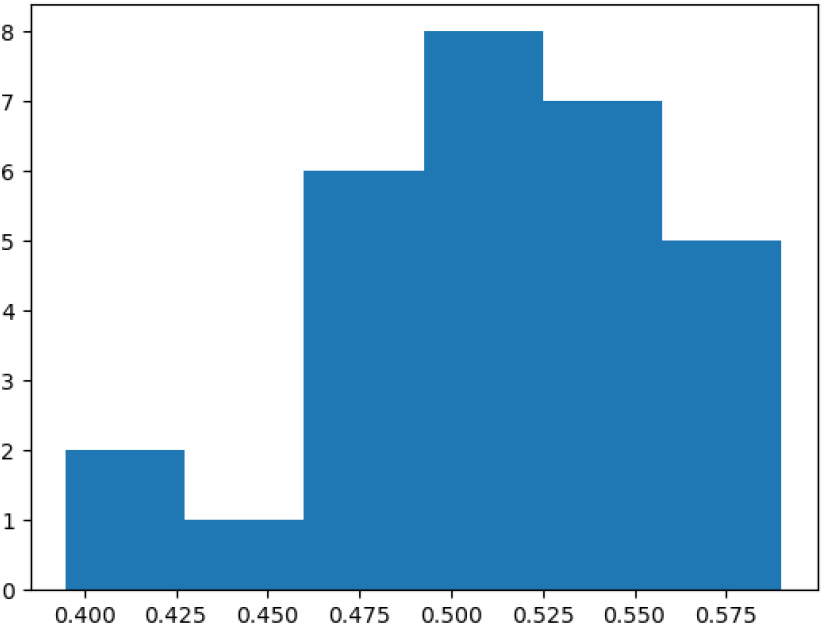
Statistics for score vector of lymphoblastic leukemia.

#### 3.3.3 Repurposing drugs and analysis

The drugs listed on the Table 5 are what the PREDR model suggested to be used for treating lymphoblastic leukemia. PREDR suggested the possibility of using a drug called Troglitazone as a treatment for the disease. Troglitazone has been used as a treatment for type 2 diabetes to date. As a result of literature review on Troglitazone, it was confirmed that the drug inhibits cell growth and induces apoptosis of b cells caused by lymphoblastic leukemia[35].

**Table 5:**
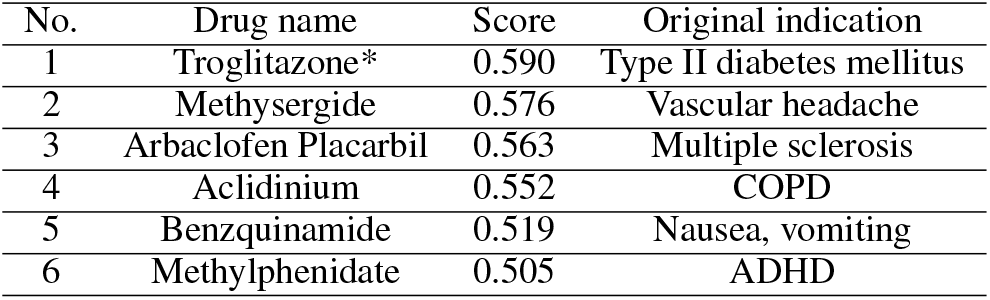
The 6 drugs are newly suggested as candidates for treating lymphoblastic leukemia by the PREDR. The top scored candidate (* marked) is used to conduct further literature studies.

We were also able to derive genes predicted to interact with Troglitazone and lymphoblastic leukemia based on the drug-gene relationship matrix and gene-disease relationship matrix generated inside the model. Since the number of genes used in the case study was 141, the interaction scores between lymphoblastic leukemia disease, each 141 genes, and the drug Troglitazone were confirmed. As for the gene-disease interaction matrix, since there was only one disease originally provided as an input, the corresponding matrix was used as it is. For the drug-gene interaction matrix, the vector of the row that mapped to the Troglitazone drug was extracted and used from a matrix of 29×141 for a total of 29 drugs. A sigmoid function was applied to each element in the obtained intermediate drug-gene and gene-disease interaction vector to calculate a score for each intermediate interaction. From the calculated score vector, 18 genes with scores greater than 0.5 in both drug-gene and gene-disease interactions were extracted. Table 6 shows the list of 18 derived genes. Literature study was additionally conducted for the four genes marked with * to investigate whether the gene actually mediates between the Troglitazone drug and lymphoblastic leukemia disease.

**Table 6:** The 18 genes listed are expected to interact to both troglitazone and lymphoblastic leukemia. “*” marked genes were additionally verified through literature study whether they mediate between Troglitazone and lymphoblastic leukemia disease.

| No. | Gene symbol | Gene name |
| --- | --- | --- |
| 1 | ANXA1 | Annexin A1 |
| 2 | BLMH | Bleomycin hydrolase |
| 3 | CACNA1S | Calcium voltage-gated channel subunit alpha 1 S |
| 4 | CDA | Cytidine deaminase |
| 5 | CYP2A6 | Cytochrome P450 family 2 subfamily A member 6 |
| 6 | CYP2B6* | Cytochrome P450 family 2 subfamily B member 6 |
| 7 | CYP2C19* | Cytochrome P450 family 2 subfamily C member 19 |
| 8 | EPHA2 | EPH receptor A2 |
| 9 | ESRRA* | Estrogen related receptor alpha |
| 10 | FPGS | Folylpolyglutamate synthase |
| 11 | GABRA6 | Gamma-aminobutyric acid type A receptor subunit alpha 6 |
| 12 | GABRB3 | Gamma-aminobutyric acid type A receptor subunit beta 3 |
| 13 | GABRG2 | Gamma-aminobutyric acid type A receptor subunit gamma 2 |
| 14 | KIT | KIT proto-oncogene, receptor tyrosine kinase |
| 15 | SERPINE1* | Serpin family E member 1 |
| 16 | RRM1 | Ribonucleotide reductase catalytic subunit M1 |
| 17 | TLR4 | Toll like receptor 4 |
| 18 | VEGFA | Vascular endothelial growth factor A |

**Table 7:** The genes listed are demonstrated the connection between troglitazone and lymphoblastic leukemia by literature studies.

| No. | Gene symbol | Relation to troglitazone | Relation to lymphoblastic leukemia |
| --- | --- | --- | --- |
| 1 | CYP2B6 | Inhibitor | Polymorphism |
| 2 | CYP2C19 | Inhibitor | Polymorphism |
| 3 | ESRRA | Inhibitor | Overexpression |
| 4 | SERPINE1 | Inhibitor | Polymorphism |

It was confirmed that 4 of the actually derived genes had an interaction between Troglitazone and each gene and lymphoblastic leukemia based on the database. It was confirmed through the DrugBank database that Troglitazone acts as an inhibitor for all four genes, and through a literature review[24], it was confirmed that each gene also acts against lymphoblastic leukemia[36–39].

## 4 Discussion

The proposed PREDR model induces the drug-disease interaction through meta-path reconstruction by merging several relations, thereby performing the drug repurposing task in a novel way that has not been attempted before. As described in Chapter 3, PREDR performed well with other drug repurposing models and showed that it could be used to explain the mechanism of drug action that other models could not. In addition to the various functions shown above, PREDR has several characteristics that have not been demonstrated experimentally, but are expected to have. We anticipate that we will be able to develop a model that performs the drug repurposing task better through additional research in the scalability of the model and the input data provided.

### 4.1 Model extensibility

In this study, the meta-path was reconstructed using only the bipartite relation of drug-gene and gene-disease. If additional proteins and bipartite relations and homogeneous relations for side effects are added, drug-disease relations for more complex meta-paths can be derived. However, there are some limitations. First, the final relation matrix obtained by multiplying the relation matrices described above represents the number of paths that can go from each source node to the destination node. (This applies only when each relation matrix is in the form of a binary adjacency matrix consisting only of 0 and 1.) A fatal problem arises here. If the network is basically dense, the connectivity from the source node to the destination node is very large, so the final matrix The number of each component gradually increases. However, in the drug repurposing task, the label is defined only as 0 or 1 because the binary classification task of whether or not there is an indication of a drug for the disease is the main one. At this time, if each component of the matrix is more than 1 in order to compare the final matrix and the label - so if more than one pathway from a specific drug to a specific disease emerges - the output should be given as 1. In that case, both a relation with one branch of the actual path and a relation with a very large number of branches are considered equally, which can negatively affect loss calculation and backpropagation. Therefore, the proposed PREDR model is expected to perform better as the given network is sparse. For this reason, if all the modules that consider various interactions are implemented and applied to the model, the aforementioned final matrix is unavoidably enlarged, so it seems that additional efforts are needed to solve this problem.

### 4.2 Representativeness of feature vectors

The proposed PREDR model can be trained without prior knowledge of each bipartite relation. This is because the relation is determined directly through the two feature vectors. However, since the model indirectly learns through the indirect task of drug repurposing after combining the information of the relation determined in each module through the product operation without directly comparing it with the golden standard, it is difficult to determine whether each relation information really reflects the biological relation. It is difficult to provide clear insight. However, since the weights used in the model are all used as learning parameters for extracting the corresponding relations, it can be expected that there will be no significant disruption to learning if the provided feature vectors reflect sufficient information about each biological entity. In this study, the dimension of the feature vector of each biological entity was limited to 5 dimensions. This is because the information that can be derived from disease is limited to 5 dimensions, and if the dimension is arbitrarily expanded, there is a possibility of falling into the curse of dimension problem. In addition, if the dimension of the feature vector of each entity is different, there is a possibility that the importance of biological entities may be considered differently in learning, so the dimension of the disease feature vector with the minimum dimension was unified. However, it is acknowledged that it may be unreasonable to evaluate the distance of the characteristics of a specific drug, gene or disease only with 5-dimensional values. Therefore, I guarantee that more accurate results can be obtained if the feature vector is derived in a more sophisticated way and used for learning to judge the relation.

## 5 Conclusion

This study is significant in that it overcame several limitations that may occur in general graph reconstruction and repurposing drugs based on drug-disease relationship information based on new meta-path reconstruction. The proposed model, PREDR, was able to confirm that it performed well with other drug repositioning models, and also has the ability to explain the reaction mechanism based on the meta-pathway for the reconstituted drug. By adding modules to the model, we expect to be able to understand drug-disease relationships for more diverse meta-pathways.

